# Structural probing of Hsp26 activation and client binding by quantitative cross-linking mass spectrometry

**DOI:** 10.1101/2021.06.06.447241

**Authors:** Julius Fürsch, Carsten Voormann, Kai-Michael Kammer, Florian Stengel

## Abstract

Small heat-shock proteins (sHSP) are important members of the cellular stress response in all species. Their best described function is the binding of early unfolding states and the resulting prevention of protein aggregation. Most sHSPs exist as oligomers but vary in size and subunit organization. Many sHSPs exist as a polydisperse composition of oligomers which undergoes changes in subunit composition, folding status and relative distribution upon heat activation. To date only an incomplete picture of the mechanism of sHSP activation exists and in particularly the molecular basis of how sHSPs bind client proteins and mediate client specificity is not fully understood. In this study we have applied cross-linking mass spectrometry (XL-MS) to obtain detailed structural information on sHSP activation and client binding for yeast Hsp26. Our cross-linking data reveals the middle domain of Hsp26 as client-independent interface in multiple Hsp26::client complexes and indicates that client-specificity is likely mediated via additional binding sites within its αCD and CTE. Our quantitative XL-MS data underpins the middle domain as the main driver of heat-induced activation and client binding but shows that global rearrangements spanning all domains of Hsp26 are taking place simultaneously. We also investigated a Hsp26::client complex in the presence of Ssa1 (Hsp70) and Ydj1(Hsp40) at the initial stage of refolding and observe that the interaction between refolding chaperones is altered by the presence of a client protein, pointing to a mechanism where interaction of Ydj1 with the HSP::client complex initiates assembly of the active refolding machinery.

## Introduction

Correct folding is critical for the function of proteins. sHSPs protect non-native proteins from irreversible aggregation by binding early unfolding states and holding them in a refolding competent state (1, 2).This allows for spontaneous folding under physiological conditions or refolding once the client protein is transferred to other ATP-consuming chaperones (3-5). sHSPS are highly conserved in all species, but vary in their size, ranging between 15 to 40 kDa. Some sHSPs are ubiquitously found and others are expressed tissue-specifically (6). sHSPs consist of a highly conserved alpha-crystallin domain (αCD), a moderately conserved C-terminal extension (CTE) and a largely unfolded N-terminal domain (NTD) (7-9). One common structural and functional feature of nearly all sHSPs is the formation of oligomers of various size (10, 11). While the αCD mediates the formation of a dimeric building block, the NTD and CTE are crucial for oligomerization (12-14). Under heat stress sHSPs switch into an activated form with increased client binding affinity and chaperone activity (15, 16). Following client binding even larger sHSP::client complexes are formed (11, 17, 18). Most of the existing literature points to the NTD as the domain that is mainly involved in client binding (19-21), even though both CTE and αCD have also been implicated in client binding and mediating client specificity (21-24). It has also been shown that the involvement of some domains appears to be client-specific and particularly the relevance of the αCD and CTE domains seem to be dependent on the nature of the client protein involved (25-27). In summary, the molecular basis of how different sHSPs bind various substrates is still not fully understood and the literature points towards a diverse binding mechanism depending on the specific client protein and sHSP involved.

The yeast genome codes for only two sHSPs, Hsp42 and Hsp26. While Hsp42 is a general and promiscuous chaperone which is responsible for protein homeostasis (28), Hsp26 is only expressed under stress conditions (29), known to interact with client proteins after heat-activation and to mediate refolding with the support of other chaperones (17, 23). In contrast to other sHSPs, Hsp26 has a remarkably long NTD (amino acids (AA): 1-95)) which is further subdivided into a N-terminal part (AA: 1-30) and a middle domain (MD, AA: 31-95) (30) that undergoes a structural rearrangement on the level of its tertiary structure upon heat activation (31). In *Saccharomyces cerevisiae* the Hsp40/Hsp70 (Ydj1/Ssa1) system acts together with Hsp104 to refold partially unfolded client proteins and this refolding activity is further enhanced when unfolded client proteins are previously bound by Hsp26 (32-34). While the function of all chaperones involved in refolding is well described, hardly any structural data of the refolding complex itself or single protein-protein interactions (PPIs) between chaperones and their client proteins within a refolding complex are available to date. One promising approach for addressing PPIs is based on the rapidly evolving technology of crosslinking coupled to mass spectrometry (XL-MS). The general approach of XL-MS is to introduce covalent bonds between proximal functional groups of proteins or protein complexes in their native environment by crosslinking reagents. The actual crosslinking sites are subsequently identified by MS and reflect the spatial proximity of the respective proteins or subunits in a complex. XL-MS provides a wealth of information on the connectivity, interaction, and relative orientation of subunits within a complex and contains spatial information, though at relatively low resolution. Relative changes in crosslinking can additionally be probed by quantitative XL-MS (qXL-MS) and we and others could show that qXL-MS can provide a structural understanding of protein dynamics (35-38).

In this work we applied XL-MS and quantitative XL-MS to different Hsp26::client complexes using three well characterized model client proteins (luciferase, malate dehydrogenase (MDH) and glutamate dehydrogenase(GDH)) as examples. We have also applied quantitative XL-MS over various time-points during heat-activation to follow structural rearrangements within Hsp26 when switching from its inactive to its active state and to monitor intramolecular changes in the overall cross-linking pattern upon client binding. Our data identify the middle domain of Hsp26 as the main driver of heat-induced activation and client binding but demonstrate additional involvement of the αCD and CTD. Finally, we have investigated Hsp26 with and without client also in the presence of the refolding chaperones Ssa1 (Hsp70) and Ydj1(Hsp40) at the initial stage of refolding and observe that the interaction of these chaperones is altered by the presence of a Hsp26::client complex, indicating that the interaction of Ydj1 with the Hsp26:client complex initiates the assembly of the active refolding machinery.

In summary, we have applied XL-MS and time-resolved qXL-MS in order to deepen our understanding of the molecular basis of heat-induced activation and client binding for yeast Hsp26 and to demonstrate how quantitative XL-MS can generally be used to monitor dynamics of sHSP::client complexes.

## Results

### Hsp26 interacts with different client proteins via similar domains

Since the molecular basis of client interaction for Hsp26 is still not fully understood, we studied protein-protein interactions (PPIs) of Hsp26 with various client proteins using XL-MS. To do so, Hsp26 was recombinantly expressed in *E. coli* and purified via a Sumo-His6 tag (**Figure S1A**). After testing the activity of Hsp26 and defining suitable reaction conditions for HSP::client complex formation (Figure S1B and S1C), we cross-linked Hsp26 in the presence of its known model client proteins luciferase, MDH and GDH at 30°C and under heat-shock conditions (**Figure 1** and **S2** and **Supplementary Table 1**). Cross-linking was performed by addition of deuterated disuccinimidyl suberate (DSS) and under an optimized regime using shorter cross-linking times than conventionally used (39) in order to ensure that the HSP: client complexes did not aggregate as response to longer cross-linking times (*see* methods for details).

**Figure 1:**
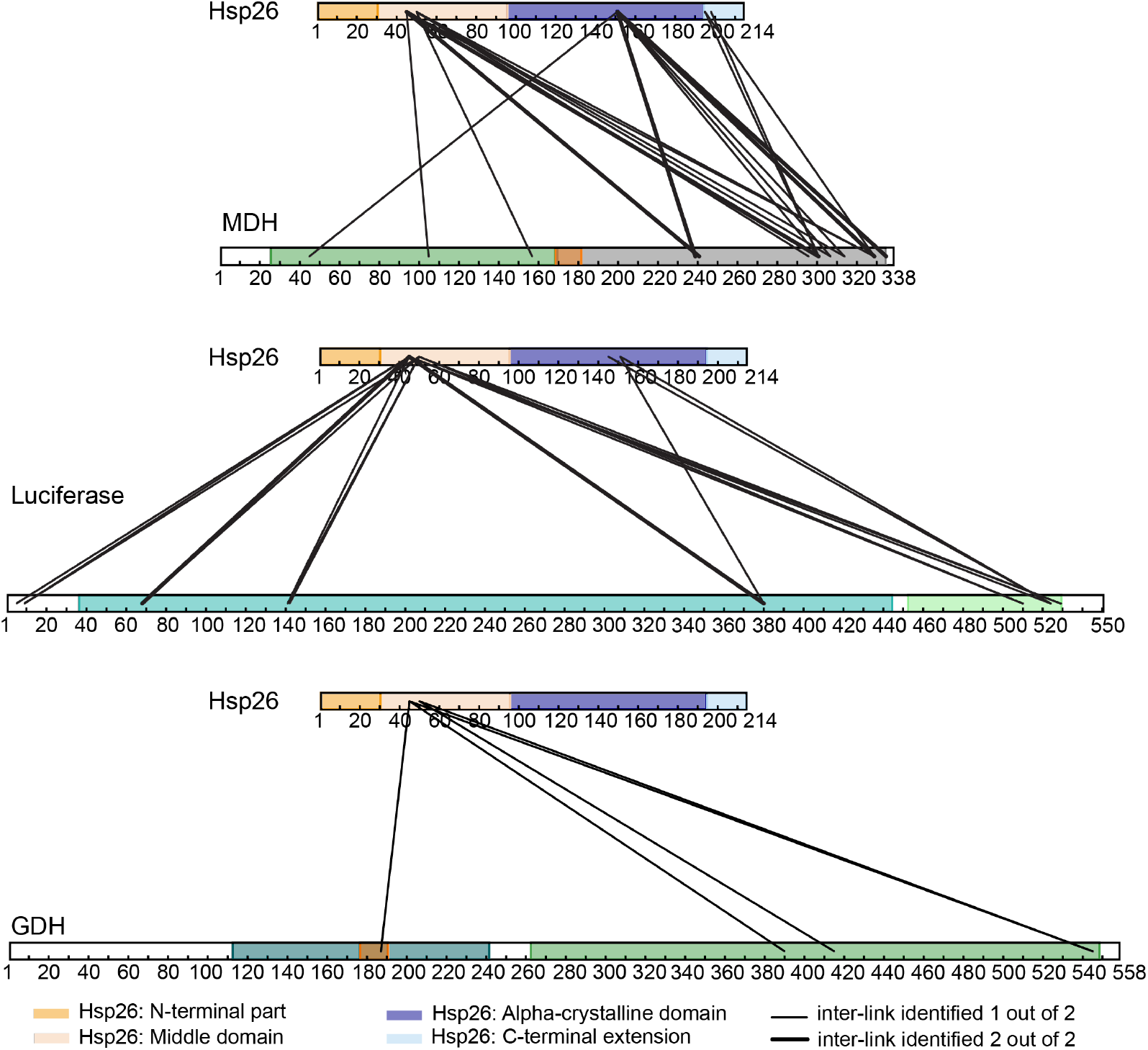
Interaction sites for three different Hsp26::client complexes identified by XL-MS. Hsp26 was incubated in the presence of the three client pro-teins MDH (top), luciferase (middle) and GDH (bottom) and cross-linked under heat-shock conditions leading to full client denaturation (see Figure S1).

Our results clearly demonstrate a heat-dependent interaction, as hardly any inter-links between Hsp26 and client proteins were detected at the ambient temperature of 30°C (**Figure S2A**). Under heat-shock conditions however, we find that Hsp26 interacted with all three client proteins via two lysines located in its MD (K45, K50) (**Figure 1**). Additionally, we identified a second binding site within the αCD (involving predominantly K151 and K145) for MDH and luciferase, while MDH additionally utilized K195 and K198 within its CTE as a third binding site. Interestingly, if client to Hsp26 ratios were further increased, these additional minor binding sites with the αCD and CTE were also utilized for client binding in case of luciferase and GDH (**Figure S2B** and **S2C**). On the client-side, binding sites were predominantly located in the C-terminal regions even though all three model client proteins engaged Hsp26 via multiple contact sites distributed all over the various client proteins (**Figure 1** and **Figure S2B** and **S2C**).

Taken together our data therefore indicates that the MD of Hsp26 acts as the main site generally involved in client binding, but that additional minor binding sites within the αCD and CTE can be utilized by Hsp26 depending on the specific client involved.

### Quantitative XL-MS enables monitoring of dynamic changes of Hsp26 during heat-induced activation and client binding over time

We now wanted to see if XL-MS, and in particularly quantitative XL-MS, could be applied to monitor dynamic changes of Hsp26 during heat-induced activation and to deepen our understanding of the molecular basis that causes Hsp26 to switch from an inactive to an active state. To do so, we extended our approach of using short cross-linking times to multiple time-points during heat activation and client binding and cross-linked aliquots of Hsp26 alone and in the presence of a client protein, both under ambient temperatures of 30°C and under heat-shock inducing conditions (**Figure 2A, Figure S3, Supplementary Data 2** and *see* methods for details).

**Figure 2:**
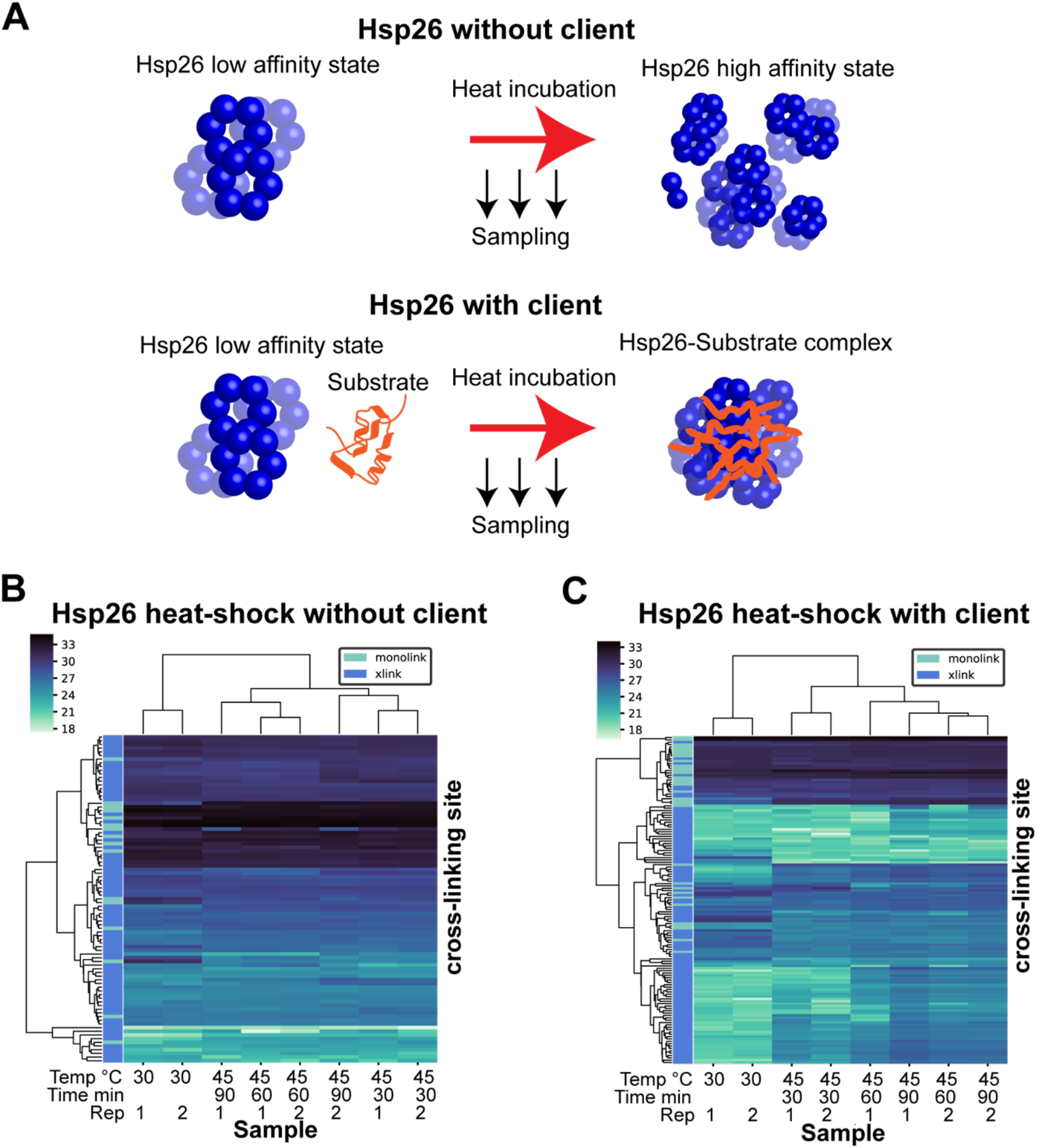
Time-resolved quantitative cross-linking mass spectrometry of Hsp26 and Hsp26:client complexes. (**a**) Schematic illustration of the experimental design. Hsp26 was heat-incubated in absence and presence of a client protein. Aliquots were taken at 30°C and at defined timepoints during heating, cross-linked with DSS (H12/D12) for 5 min and analyzed by quantitative cross-linking mass spectrometry. (**b**) Hierarchical clustering of cross-link and monolink quantities of Hsp26 in a time course experiment reveals changes in the overall cross-linking pattern. The experiment was carried out as described in (a) using an incubation temperature of 45°C. Samples were taken at indicated timepoints (30 min, 60 min and 90 min). (**c**) Hierarchical clustering of cross-link and mono-link quantities of Hsp26 and MDH.

Hierarchical clustering of cross-link quantities using in-house scripts (*see* methods for details) allowed us to quickly judge similarities and differences in cross-linking patterns on a global scale. **Figure 2B** shows exemplarily one of the time-course experiments in absence and presence of the client protein MDH. The samples processed under ambient temperatures cluster separately from the samples processed under heat-shock conditions - independent of client protein present - while the various heat-shock clusters show high intrinsic similarities. This global representation of our data by means of hierarchical clustering clearly shows that qXL-MS has the ability to detect differences in cross-linking patterns in a time-resolved manner and thus to monitor dynamic changes of Hsp26 during heat-induced activation and client binding.

### qXL-MS identifies middle domain of Hsp26 as the main driver of heat-induced activation and client binding

In a next step we applied these settings to study activation and client binding for Hsp26. **Figure 3** and **Figure S4** show changes in the cross-linking pattern of Hsp26 during heat-induced activation in the absence or presence of the client protein MDH (*see* also **Supplementary Table 2**). While **Figure S4** focusses on the level of the single cross-linking site, **Figure 3** takes this data as an input to allow for more generalized statements.

**Figure 3.**
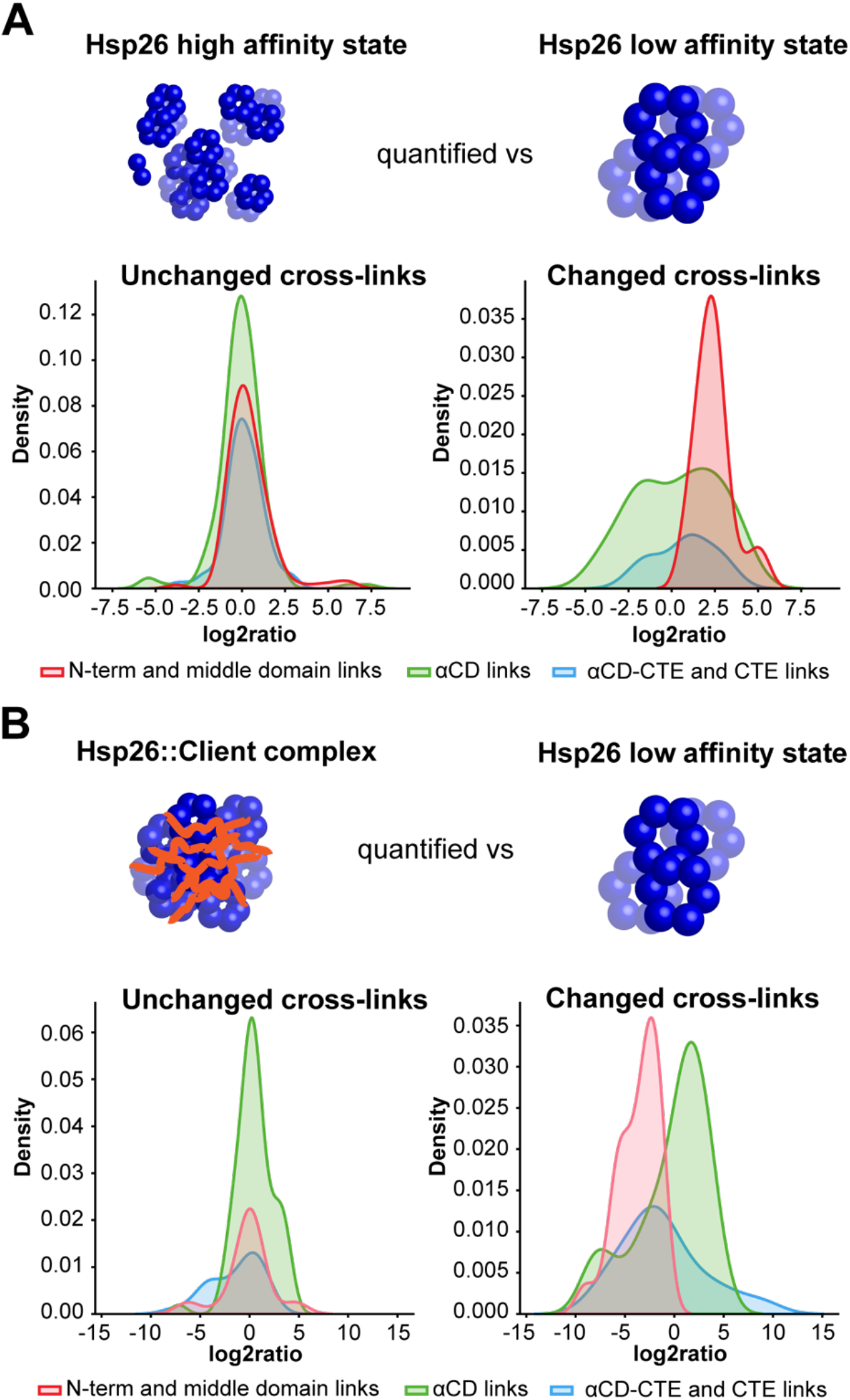
Domain specific changes of Hsp26 during heat-induced activation and client binding. Shown are domain specific changes in Hsp26 overall cross-link patterns during heat-induced activation in the absence (**a**) or presence (**b**) of the client MDH. Heat-shock conditions (high affinity state) were quantified versus ambient temperature at 30°C (low affinity state) and relative changes in cross-link quantity were calculated for each unique cross-linking site, merged, and grouped into non-significant and significant changes and plotted for each of the domain groups separately (*see* Figure S4 and methods for details).

On the level of the single cross-linking site, we observed significant and reproducible changes in the cross-link abundances in all domains of Hsp26 during heat activation of Hsp26, both for Hsp26 on its own and in the presence of client protein.

In case of Hsp26 alone, we detect multiple cross-links emanating from the MD into other domains and find in particularly a cluster of links bridging the MD and the αCD that were significantly increased upon heat shock (**Figure S4A**). If we take a more general look, we find our notion confirmed that most notably cross-links involving the NTD and especially the MD change during heat activation (**Figure 3A)**, indicative of structural changes (36) and pointing towards a crucial role of the MD during the transition of Hsp26 into its high affinity state. In the presence of client protein, we also observed significant changes of cross-links between the MD and the αCD (**Figure S4B**). In this case however, these cross-links were significantly decreased, which stands in clear contrast to the data we obtained for Hsp26 on its own. This data could in principle be explained by the binding of the client protein itself to the MD and would lend additional support to the important role of the MD in client binding. As in the Hsp26 alone samples, the αCD showed a mixed population of increased and decreased links in presence of client protein, while αCD - CTE cross-links in presence of a client protein were significantly decreased, indicating again a role in client binding (**Figure S4B**). Our global view again supports the dominant role of the NTD, and in particularly the MD, in client binding, even though the effect is less pronounced than in the case of Hsp26 on its own and supports additional involvement of the αCD and CTE (**Figure 3B**). Interestingly, if we take a closer look at selected cross-linking sites and follow their changes in abundance in a more time-resolved manner during heat-induced activation and client binding, we observe that many changes take place already during the first minutes of incubation under heat shock temperatures (**Figure S4C**).

Taken together, we find that the MD exhibits the strongest changes in the transition of Hsp26 from the low to its high affinity state and during formation of the Hsp26::client complex.

### Interactions of Hsp26 and client protein with the refolding system Ydj1:Ssa1

While the functional role of Hsp40 (Ydj1) and Hsp70 (Ssa1) as refolding system in cooperation with sHSPs is well described (32, 33), there is hardly any structural information on the interaction of the refolding system Ydj1::Ssa1 with Hsp26 and its clients. We expressed and purified Ydj1 and Ssa1 recombinantly in *E*.*coli* to obtain structural information on PPIs of the Hsp26::Ydj1::Ssa1 refolding machinery and client protein (**Figure S5A**). We then performed a refolding assay to test for refolding activity and observed refolding of luciferase by Ydj1::Ssa1 as previously described (33, 34) (**Figure S5B**).

In a next step we then cross-linked Hsp26 and the refolding chaperones Ssa1 (Hsp70) and Ydj1(Hsp40) in the presence or absence of luciferase. To mimic the cellular system as closely as possible, Hsp26 was first heat activated in absence and presence of luciferase and subsequently allowed to cool down, before Ydj1 and Ssa1 were added to the Hsp26::client complex and crosslinking of the refolding complex (**Figure 4** and **Supplementary Table 3** and *see* methods). **Figure 4A** and **4B** show the overall inter-protein cross-linking patterns of the refolding complex in the presence and absence of client protein luciferase. While the refolders Ydj1 and Ssa1 interact widely with each other without client protein present, this picture changes with the arrival of the Hsp26::client complex. Now the interprotein cross-links between Ydj1 and Ssa1 become significantly less pronounced, and the Hsp26::luciferase complex engages in multiple interprotein cross-links with Ydj1.

**Figure 4.**
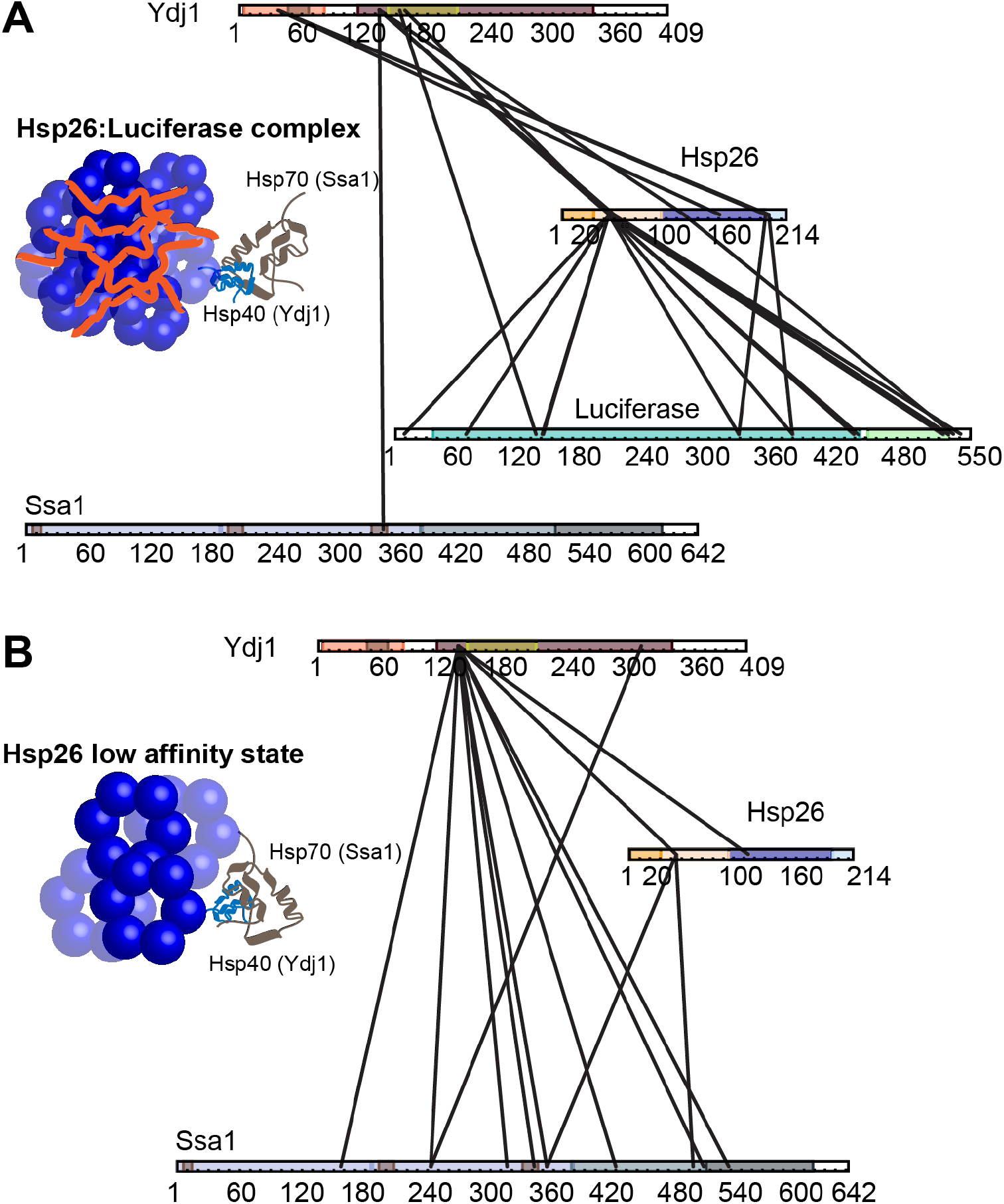
Cross-linking of a Hsp26::client complex in the presence of Ydj1/Ssa1. (**a**) Inter-protein cross-links within the Hsp26::luciferase::Ydj1::Ssa1 refolding complex. (**b**) Inter-protein cross-links of Hsp26, Ydj1 and Ssa1 in absence of a client protein.

These results suggest a mechanism where first Ydj1 interacts directly with the Hsp26::client complex in order to pass the denatured client protein on to Ssa1 for refolding and is therefore responsible for the assembly of the active refolding complex.

## Discussion

In this work we applied XL-MS to different Hsp26::client complexes using three well characterized model clients. While we identified binding sites within all three domains of Hsp26, which were similar for all investigated client proteins, the MD showed by far the most inter-links. This is in line with previous studies which could demonstrate that all domains of a sHSP can play a role for chaperone activity (22, 23, 40) but that the MD domain is essential for client binding in Hsp26 (30). Moreover, our data shows that the inter-links involving the MD were formed already at lower amounts of denatured client protein present, indicating that this domain comprises the main client binding site, independent of the client protein involved. In contrast, the degree to which additional binding sites within the αCD and CTE were utilized within Hsp26 appeared to largely depend on both the specific client involved and the amount of client protein present, as suggested previously (21, 25-27). Our data is therefore in line with an interpretation where initial client engagement via the NTD of sHSPs, potentially coinciding with a conformational change within the NTD (21, 26), sets the stage for additional binding events involving additional domains depending on the specific needs and the client protein involved. So far, it is still not fully understood how sHSPs mediate their broad client specificity (41) and it remains an open question if client specificity is achieved mainly due to their polydispersity or because HSPs are able to recognize and exploit various binding interfaces. In our study we found that Hsp26 uses very similar regions for its interaction with three model client proteins. On the client-side, binding sites were predominantly located in their C-terminal regions, even though we observed that all client proteins engaged Hsp26 via multiple regions distributed over the respective client protein. This points to a mechanism where client specificity is not mediated via defined interfaces with distinct properties but rather to a mechanism where structural heterogeneity and oligomeric polydispersity accounts for the widespread client specificity observed for Hsp26 and HSPs in general. We and others could show previously that relative changes in cross-linking quantities as probed by quantitative XL-MS can provide a structural understanding of protein dynamics (35-38). Here we have used this approach to probe the dynamics of sHSP::client complexes. By using quantitative XL-MS and shortened cross-linking times we were able to follow the activation process of Hsp26 during heat stress and the stepwise formation of a Hsp26::client complex over time. Our data revealed significant changes in cross-linking patterns, likely associated with structural rearrangements, as response to heat-shock, in line with previous studies (17, 31, 42). While we observed changes in cross-link quantity within all domains of Hsp26, we identify the middle domain of Hsp26 as the main driver of heat-induced activation and client binding. Particularly, we find that the majority of cross-links between the MD and the αCD and the MD and the CTE were increased in the absence of client protein, a process which was virtually reversed in the presence of client protein. The role of the MD for the activation of Hsp26 had been demonstrated previously in a study which identified the MD as a “thermosensor” that undergoes an intrinsic conformational change upon heat activation (31). In our data the αCD itself also exhibited strong changes in the absence and presence of client. However, in this case we observed a mixed effect consisting of increased and decreased cross-link abundances which were difficult to interpret and are most likely also influenced by neighbouring regions. In addition to the above-described changes in crosslink abundances in connection with the MD, we observed a decrease of αCD-CTE cross-links in the presence of a client protein. It has been reported that the CTE participates in the dimerization of the αCD (43), plays a role in oligomerization (31), is in close spatial proximity to the αCD (12) and plays a role in client interaction (23, 25, 26). Our data therefore indicates that αCD and CTE may act together in client binding and that the MD is essential for both heat induced activation of Hsp26 and client binding.

Finally, we have investigated Hsp26 also in the presence of the refolding chaperones Ssa1 (Hsp70) and Ydj1(Hsp40) and see that interaction between these refolding chaperones is altered by the presence of a Hsp:26::client protein complex. Type A Hsp40s as Ydj1 comprise a highly conserved J-domain, a ZnF and a C-terminal domain. Both the ZnF and C-terminal domain have been shown to play an important role in client binding (44, 45). In their client-bound state Hsp40s stimulate the ATPase activity of Hsp70s via a functional interaction of their J-domain and the Hsp70 nucleotide-binding domain (NBD) (46), thereby presenting the unfolded client to the substrate-binding domain (SBD) of the respective Hsp70 chaperone. Recently, it was shown that the J-domain can also interact with the SBD directly (47) and that in case of Ydj1 the client is bound by a hydrophobic pocket within its C-terminal domain comprising amino acids I116, L135, L137, L216 and F249 (48). Mutational studies have revealed that both ZnF motifs of Ydj1 are important to assist in Hsp70 mediated refolding, but that only the first motif (C143,C146 and C201,C204) plays a role in client binding while the second one (C159,C162 and C185,C188) appears to be important for the interaction with Hsp70 (49). In our data we have detected inter-protein cross-links between lysines K134 and K158 of Ydj1 with lysines K45 and K50 located in the MD of Hsp26. As K158 is neighbouring to the second ZnF motif in Ydj1 and as K134 is spatially close to its client binding pocket, our data may hint at a mechanism where the unfolded client protein is taken over by the Ydj1 binding pocket from the Hsp26::client complex by interacting with the MD of Hsp26 domain, potentially with assistance of the ZnF. We also detected additional inter-links between the J-domain of Ydj1 and the αCD and CTE of Hsp26 that are exclusive to the client-bound state. This indicates that Ydj1 may engage in a second interface with the Hsp26::client complex independent of the MD domain of Hsp26 and gives further support to the notion that the J-domain may have additional roles to its well described stimulation of Hsp70’s ATPase activity (46), even though future investigations will be needed to resolve this question.

Apart from the interaction of Ydj1 with the Hsp26::client complex, our most striking result is the dynamics of the interaction between Ydj1 and Ssa1. While the refolders interact widely with each other without client protein present, this picture changes dramatically with the arrival of the Hsp26::client complex. Now the interprotein cross-links between Ydj1 and Ssa1 become significantly less pronounced, and the Hsp26::luciferase complex engages in multiple interprotein cross-links with Ydj1. Taken together our data suggests that binding of unfolded client protein by Hsp26 and subsequent interaction of Ydj (Hsp40) and the Hsp26::client complex initiates the assembly of the active refolding machinery. In summary, we have applied XL-MS and time-resolved qXL-MS in order to deepen our understanding of the molecular basis of heat-induced activation and client binding for yeast Hsp26 and to demonstrate how quantitative XL-MS can generally be used to monitor dynamics of sHSP::client complexes and to structurally probe the eukaryotic chaperone machinery.

## Material and Methods

### Expression and purification of recombinant proteins

Chaperones were expressed in E. coli (Rosetta2 (DE3) pET plasmid expressing His6-Sumo-Hsp26/Ydj1/Ssa1). In short, a pre-culture was inoculated by a single-cell colony and used to inoculate a second overnight culture. The main culture was inoculated with a starting OD600 of 0.1 and grown at 30°C and 120 rpm. At an OD600 of 0.6 the recombinant protein expression was induced by 1 mM IPTG and overexpression was performed for 4 hours. The cells were harvested by centrifugation at 4500xg for 20 min and the pellet was resuspended in lysis buffer (40 mM Hepes (pH 7.4), 500 mM NaCl (Hsp26), 150 mM NaCl (Ydj1 and Ssa1), 20 mM Imidazole, 1 mM TCEP) and centrifuged a second time before the pellet was snap frozen in liquid nitrogen and stored at −80°C. The cell pellet was thawed in lysis buffer at 37°C for 15 min. The lysis buffer was supplemented freshly by a spatula tip of magnesium chloride, DNAse I and protease inhibitors Aprotinin/Leupeptiden (1 mg/L) and Pefabloc® (100 µM). For complete cell lysis the cells were sonicated with a micro-tip at 40 % amplitude 3 times for 10 seconds. After that the cell lysate was clarified by centrifugation at 19500xg for 40 min. The supernatant was transferred to a superloop for purification by FPLC. A glass column (YMC) was self-packed with Ni-IDA Protino (Machery-Nagel) beads and a flow rate of 0.5 mL/min was used. Cell lysate was loaded onto the column with lysis buffer as washing buffer. Elution was done with lysis buffer always containing 500 mM NaCl and 1 M Imidazol. A gradient from 0 to 100 % elution buffer was run for 20 mL. Target elution fractions were identified by SDS-PAGE and pooled. In house provided Ulp1 was added in a ratio of roughly 1:200 (w/w) to the pooled elution fractions to cleave off the Sumo-His6 Tag. The pooled elution fractions were dialyzed with a 3kDa MWCO dialysis tubing (snakeskin, Thermo Scientific) in 4 L dialysis buffer (40 mM Hepes (pH 7.4), 150 mM NaCl, 1 mM β-ME) overnight to reduce the imidazol concentration and to allow tag cleavage. The next day tag removal was either done by a second Ni-IDA affinity purification (same buffers, 1 mL/min flow rate, 5 mL imidazol gradient) or by size exclusion chromatography (SEC) using a Superdex® 75 10/300 GL (GE Healthcare) with a flow rate of 0.5 mL/min and dialysis buffer as running buffer. Target proteins were always base line separated from the tag since all chaperones eluted in the void volume. The target fractions were identified by SDS-PAGE, pooled and concentrated to roughly 0.5 – 1 mL by ultrafiltration spin units Amicon® Ultra–15 10 kDa MWCO (Merck-Millipore). Buffer was exchanged against storage buffer (40 mM Hepes (pH 7.4), 50 mM NaCl (Hsp26), 150 mM NaCl (Ydj1 and Ssa1), 1 mM EDTA, 10% Glycerol) within the same spin unit. Purified and concentrated protein was aliquoted, snap frozen in liquid nitrogen and stored at 80°C. Protein concentration was determined by BCA as well as Bradford assay.

### Aggregation assay

To check the activity of Hsp26 to inhibit substrate aggregation, we performed an aggregation assay (often referred as light scattering). The substrates, Hsp26 alone and Hsp26 + client, respectively were diluted in heating buffer (40 mM Hepes (pH 7.4), 50 mM NaCl, 0.5 mg/mL BSA, 2 mM β-ME) and incubated in plastic cuvettes in a heated water bath, respectively. Light scattering was measured at 550 nm in a dual-beam photometer with heating buffer as reference. Light scattering was measured before heating and at indicated time points while heat-incubation. The substrates were purchased commercially. L-malate dehydrogenase (MDH, from pig heart mitochondrial, Roche), glutamate dehydrogenase (GDH, from bovine liver, Sigma Aldrich, G2501) and luciferase (from firefly, Sigma Aldrich, L9420).

### Client inactivity assay

To ensure that the substrates were fully unfolded, we checked the decrease of enzyme activity during heat-incubation. To do so, we incubated the client in 1.5 mL tubes in a heated thermomixer and measured enzyme activity in microplate format at indicated time points. The assay was necessary because cross-linking was as well carried out in the same thermomixer and conditions of the aggregation assay were not directly transferable. The substrate stock solutions were diluted in heating buffer and samples were taken while heating. 5 µL of heated substrate were mixed with 95 µL enzyme activity buffer. In case of MDH and GDH the oxidation of NADH to NAD+ was measured at 340 nm. In case of luciferase the decrease of luminescence was measured. GDH activity buffer: 85 mM Tris (pH 7.4), 0.5 mM NADH, 7.6 mM α-ketoglutarate, 125 nM GDH (final assay concentration). Luciferase activity buffer: 85 mM Tris-HCl (pH 7.4), 0.05 mM luciferin, 1 mM ATP, 0.5 mg/mL BSA, 15 mM MgCl2, 80 nM Luciferase (final assay concentration). MDH activity assay buffer: 150 mM potassium phosphate buffer (pH 7.4), 10 mM DTT, 0.5 mM oxaloacetate (fresh, on ice), 0.28 mM NADH, 5 nM MDH (final assay concentration).

### Luciferase refolding assay

The activity of Ydj1/Ssa1 was measured by a luciferase refolding assay. 80 nM luciferase (monomer) was denatured in presence of 160 nM Hsp26 (dimer) at 40°C for 40 min in heating buffer. After heating, the sample was cooled down at 30 °C for 30 min. 5 µL of the Hsp26::client complex were mixed with 90 µL refolding buffer and 5 µL of pre-mixed refolding proteins (1 µM Ssa1 monomer and 1µM Ydj1 dimer) in a micro-plate. As control the Hsp26::client complex was used in absence of refolding chaperones. Luminescence was monitored constantly without addition of fresh substrate over two hours at 30°C. Refolding buffer: 40 mM Hepes, 50 mM NaCl, 15 mM MgCl_2_, 0.05 mg/mL BSA, 1 mM ATP, 0.1 mM luciferin, 2 mM β-ME, 0.01 µM pyruvate kinase, 1.5 mM phosphoenolpyruvate

### Cross-linking coupled to Mass Spectrometry (XL-MS)

5 nmol Hsp26 (dimer) was incubated alone or in presence of substoichemtric amounts of client protein (luciferase, MDH or GDH, *see* Supplementary Data 4 for protein concentrations) in heating buffer at 30°C for 20 min or under heat-shock conditions. Cross-linking was performed with DSS at the same temperature for 1 min or 5 min with 2.7 molar excess of DSS to lysines. The reaction was stopped by snap freezing in liquid nitrogen and quenched by addition of 50 mM Ammonium Hydrogen Carbonate while thawing and incubation for 10 min at room temperature. The samples were evaporated to dryness. For cross-linking of the refolding complex, Hsp26 was heated in absence or presence of luciferase at 40°C for 40 min in heating buffer supplemented with 15 mM MgCl_2_ and subsequently cooled down for 30 min at 30°C before Ydj1/Ssa1 and 10 mM ATP were added. Cross-linked samples were processed essentially as described (50). The dried protein samples were denatured in 8 M Urea, reduced by addition of 2.5 mM TCEP at 37°C for 30 min and subsequently alkylated using 5 mM Iodoacetamide at RT for 30 min in the dark. After adding 50 mM ammonium hydrogen carbonate to a final concentration of 1 M urea, the samples were digested by addition of 2 % (w/w) trypsin (Promega) over night at 37°C. Digested peptides were separated from the solution and retained by a C18 solid phase extraction system (SepPak Vac 1cc tC18 (50 mg cartridges, Waters) and eluted in 50 % ACN, 0.1 % FA. After desalting the peptides were evaporated to dryness and stored at −20°C. Dried peptides were reconstituted in 30 % ACN, 0.1 % TFA and then separated by size exclusion chromatography on a Superdex 30 increase 3.2/300 (GE Life Science) to enrich for cross-linked peptides. Two or three early-eluting fractions were collected for MS measurement and enriched fractions were evaporated to dryness. Peptides were reconstituted in 5 % ACN, 0.1 % FA and protein concentrations were normalized on the peptide level based on the A215 nm absorbance to ensure equal amounts of peptides for MS measurement. The peptides were separated on a PepMap C18 2µM, 50 µM x 150 mm (Thermo Fisher) using a gradient of 5 to 35 % ACN for 45 min. MS measurement was performed on an Orbitrap Fusion Tribrid mass spectrometer (Thermo Scientific) in data dependent acquisition mode with a cycle time of 3 s. The full scan was done in the Orbitrap with a resolution of 120000, a scan range of 400-1500 m/z, AGC Target 2.0e5 and injection time of 50 ms. Monoisotopic precursor selection and dynamic exclusion was used for precursor selection. Only precursor charge states of 3-8 were selected for fragmentation by collision-induced dissociation (CID) using 35 % activation energy. MS2 was carried out in the Ion Trap in normal scan range mode, AGC target 1.0e4 and injection time of 35 ms.

Data were searched using *xQuest* in ion-tag mode (50). Carbamidomethylation (+57.021 Da) was used as a static modification for cysteine. As database the sequences of the measured recombinant proteins and reversed and shuffled sequences were used for the FDR calculation by *xProphet*. Cross-links were only considered, if they were identified with a deltaS < 0.95, a minimum Id score ≥ 25 (filtering was done on the level of the unique cross-linking site) and had additionally an FDR ≤ 0.05 as calculated by *xProphet* for at least one of the replicates within a related experiment.

### Quantitative XL-MS (qXL-MS)

Initial processing of identified cross-linked peptides for quantitation was performed essentially as described (36). In short, the chromatographic peaks of identified cross-links were integrated and summed up over different peak groups for quantification by *xTract* (taking different charge states and different unique cross-linked peptides for each unique cross-linking site into account. Only high-confidence cross-links that fulfilled the above introduced criteria were selected for further quantitative analysis. The resulting *bagcontainer*.*details*.*stats*.*xls* file was used as an input for in-house scripts. The bag container contains all experimental observations on a peptide level as extracted by *xTract* (e.g. peptide mass, charge state, the extracted MS1 peak area and any violations assigned by *xTract)*. Missing observations were replaced by imputation with random values drawn from a normal distribution based on our experimental distribution. Here, the log-normal experimental distribution of measured MS1 peak areas was converted to a normal distribution by log2-conversion. Of the resulting normal distribution, the mean and standard deviations were determined. The mean is shifted downward while the width is decreased in order to obtain our distribution to draw imputed values from. This follows the same procedure and parameters as described for Perseus (51) (width: 0.3 and down shift: 1.8). Data were additionally filtered using a light-heavy filter as described (52). Since a light-heavy cross-linker at 1:1 ratio was used, we would expect that a given peptide for a specific charge state is found with nearly the same MS1 peak area for both its heavy and light form. A peptide for a given charge state therefore receives a violation (and is subsequently filtered) if its light-heavy log2ratio is smaller than 0.5 or greater than 2. Experiments were normalized by using the mean MS1 peak areas of all experiments as the reference. The ratio of all experiments compared to the reference was computed and all observed MS1 areas were multiplied by this experiment-specific ratio to receive the same mean for all experiments. In addition, replicates were normalized within each experiment. This means that the mean of each technical replicate within an experiment is shifted to the experiment mean in the same way as described for the experiment normalization. In a next step log2ratios were calculated as the difference between the log2-converted MS1 peak areas instead of the ratio. Here, the MS1 areas for each experiment are shifted into log2 scale after all summing operations but before taking any means allowing us to calculate meaningful standard deviations between biological replicates avoiding the influence by outliers in the original log-normal scale. P-value calculation was otherwise done as described (52) with one notable exception: MS1 peak areas were not split by technical replicates to avoid artificially improving p-values with increasing numbers of technical replicates. FDR values are p-values corrected for multiple testing, following the Benjamin–Hochberg procedure.

### Statistics and reproducibility

The Hsp26::client interaction experiments were carried out in 3 biological replicates for each client (two replicates are shown in Figure 1 and the third replicate with a higher client to Hsp26 ratio is additionally included in Figure S2). The qXL-MS experiment for Hsp26 on its own (Figure 2B, 3A and S4A) was carried out in 3 biologically independent sets of experiments each consisting of cross-linking duplicates (i.e. cross-linking, sample processing and LC-MS/MS was carried out separately). The qXL-MS experiment for Hsp26 in presence of MDH (Figure 2C, 3B and S4B) was carried out in two biologically independent sets of experiments each one consisting of cross-linking duplicates (*vide supra*). The refolding experiment (Figure 4A+B and Figure S5C+D) was carried out in 4 independent replicates each one consisting of cross-linking triplicates (*vide supra*). All samples were additionally measured as technical duplicates

### Visualization

The plot shown in Figure 2 is based on a customized seaborn (*version 0*.*9*.*0*) clustermap with a Canberra distance metric and uses the log2-converted MS1 areas to cluster biological replicates by similarity. The plot shown in Figure 3 is based on a seaborn distribution plot with the kind=”kde” setting and depicts the kernel density estimator (KDE) for each condition. The KDE approximates the probability function that generated the dataset. It is similar to a histogram but without defined bins; a Gaussian component is associated with each point leading to a mixture showing a smooth function. Each conditional density (represented by a colour) was scaled by the number of observations in order for the total area under all curves for each sub figure to sum up to one. The log2ratios were again calculated using our in-house quantitation script.

### Data availability

All data generated or analyzed during this study are included in this published article and its Supplementary information files. The MS raw files have been deposited to the ProteomeXchange Consortium via the PRIDE partner repository (53) with the dataset identifier PXD026244 (username: and password: NcQUa43H).

## Author Contributions

J.F. and F.S. conceived the study and experimental approach; J.F. expressed and purified proteins with help from C.V.; J.F. performed all assays and XL-MS experiments with help of C.V.; J.F., K.M.K., and F.S. analyzed the data, and J.F. and F.S. wrote the paper with input from all authors.

## Acknowledgements

This work was funded by the German Science Foundation Emmy Noether Programme to F.S. (STE 2517/1-1). F.S. also acknowledges funding from the Konstanz Research School Chemical Biology (KoRS-CB) and support from the DFG Collaborative Research Centre (SFB) 969.

## Declaration of Interests

The authors declare no competing interests.

## Supplemental Figures

**Figure S1:**
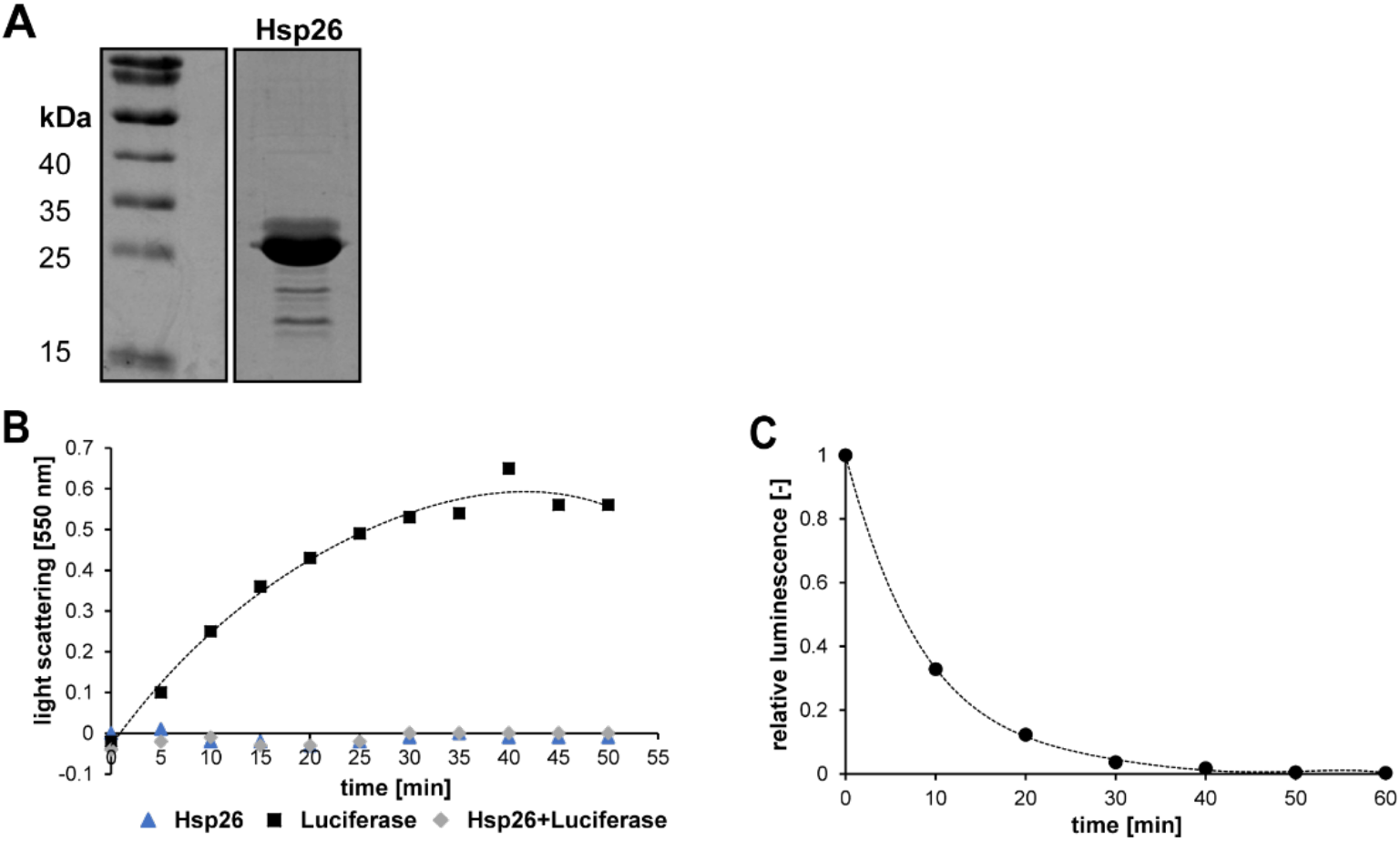
**(a)** SDS-gel of 10 µg of recombinantly expressed and purified Hsp26. **(b)** Aggregation assay of Hsp26 and luciferase. 1.25 µM luciferase (Monomer) was incubated at 40°C in absence and presence of 2.5 µM (dimer) Hsp26 and light scattering was measured at 550 nm (Hsp26 alone (blue triangles), luciferase alone (black squares) and Hsp26/luciferase (grey tilted squares)). **(c)** Inactivation assay of luciferase. 80 nM luciferase was incubated at 40°C and activity was measured at indicated time points.

**Figure S2.**
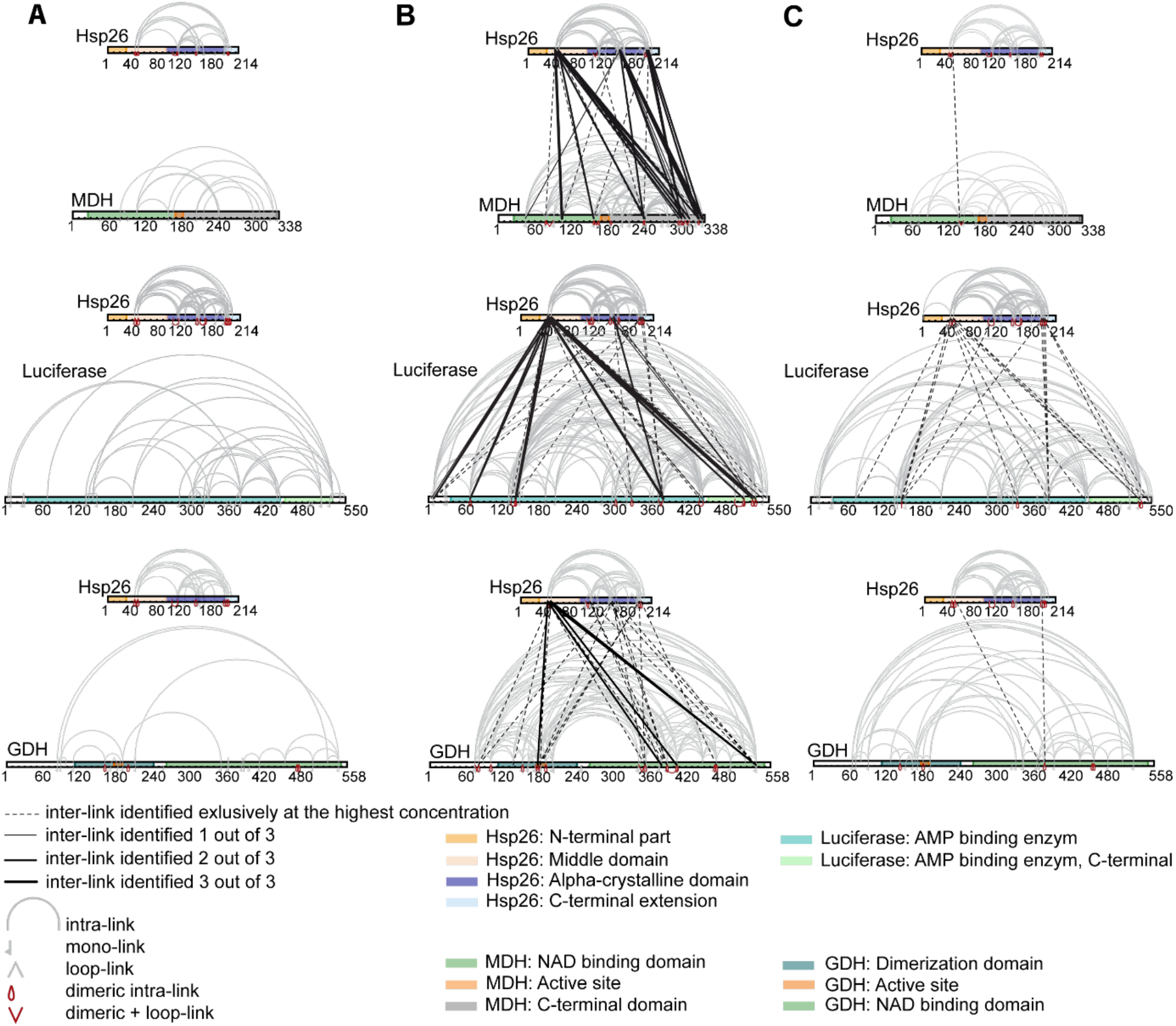
Interaction sites for three different Hsp26::client complexes identified by XL-MS. **(a)** Hsp26 was incubated in the presence of the three client proteins MDH (top), luciferase (middle) and GDH (bottom) and cross-linked at the physiological temperature of 30°C. **(b)** Hsp26 was incubated in the presence of the three client proteins MDH (top), luciferase (middle) and GDH (bottom) and cross-linked under heat-shock conditions leading to full client denaturation (*see* Figure S1) and +/-high client ratio, *see* methods and Supplementary Data 1 and 4 for details. **(c)** Hsp26 was incubated in the presence of the three client proteins MDH (top), luciferase (middle) and GDH (bottom) and cross-linked at the physiological temperature of 30°C and +/-high client ratio, *see* methods and Supplementary Data 1 and 4 for details. At this very high client concentration inter protein cross-links were also detected under physiological conditions, most likely due to protein instability (luciferase) under these conditions or due to particularly transient interactions.

**Figure S3.**
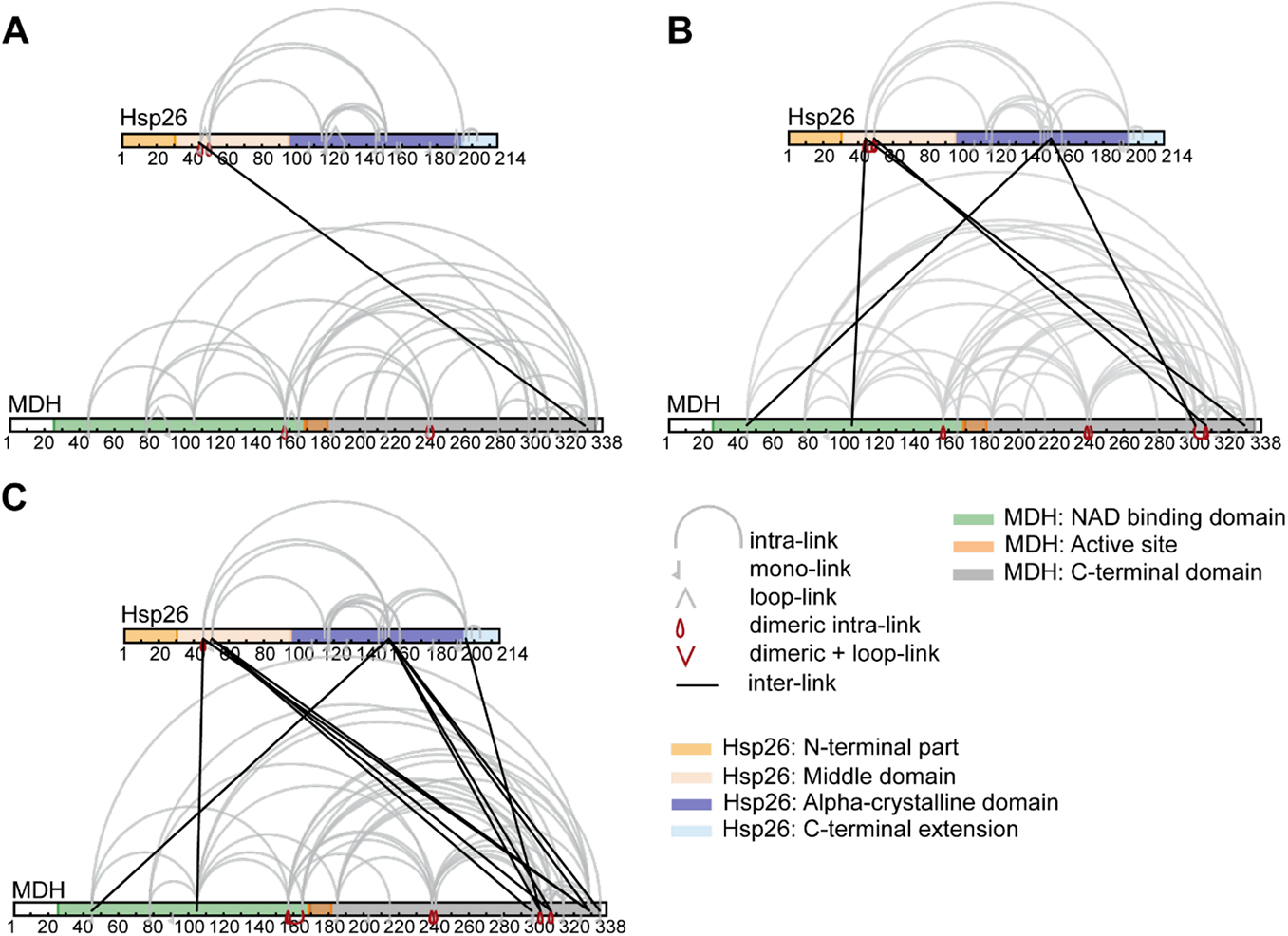
Time course experiment of Hsp26 and MDH. 5 nmol Hsp26 (dimer) were incubated with 1 nmol MDH (dimer) at 45 °C for 90 min and subsequently crosslinked with DSS (H12/D12) for 5 min at the same temperature (*see* Supplementary Data 2 and 4 for details). **(a)** Overall cross-linking pattern after 30 min. **(b)** Overall cross-linking pattern after 60 min. **(c)** Overall cross-linking pattern after 90 min. The data indicates that client binding via the MD takes place already at the earliest time point investigated when client protein has only started to denature (a), whereas binding via the αCD takes place at later timepoints (b) and interaction via the CTE only occurs at the final stage of heat-induced client denaturation (c). These results therefore support the notion that the MD is the primary client binding site and that interaction via the αCD and CTE has a supportive function.

**Figure S4.**
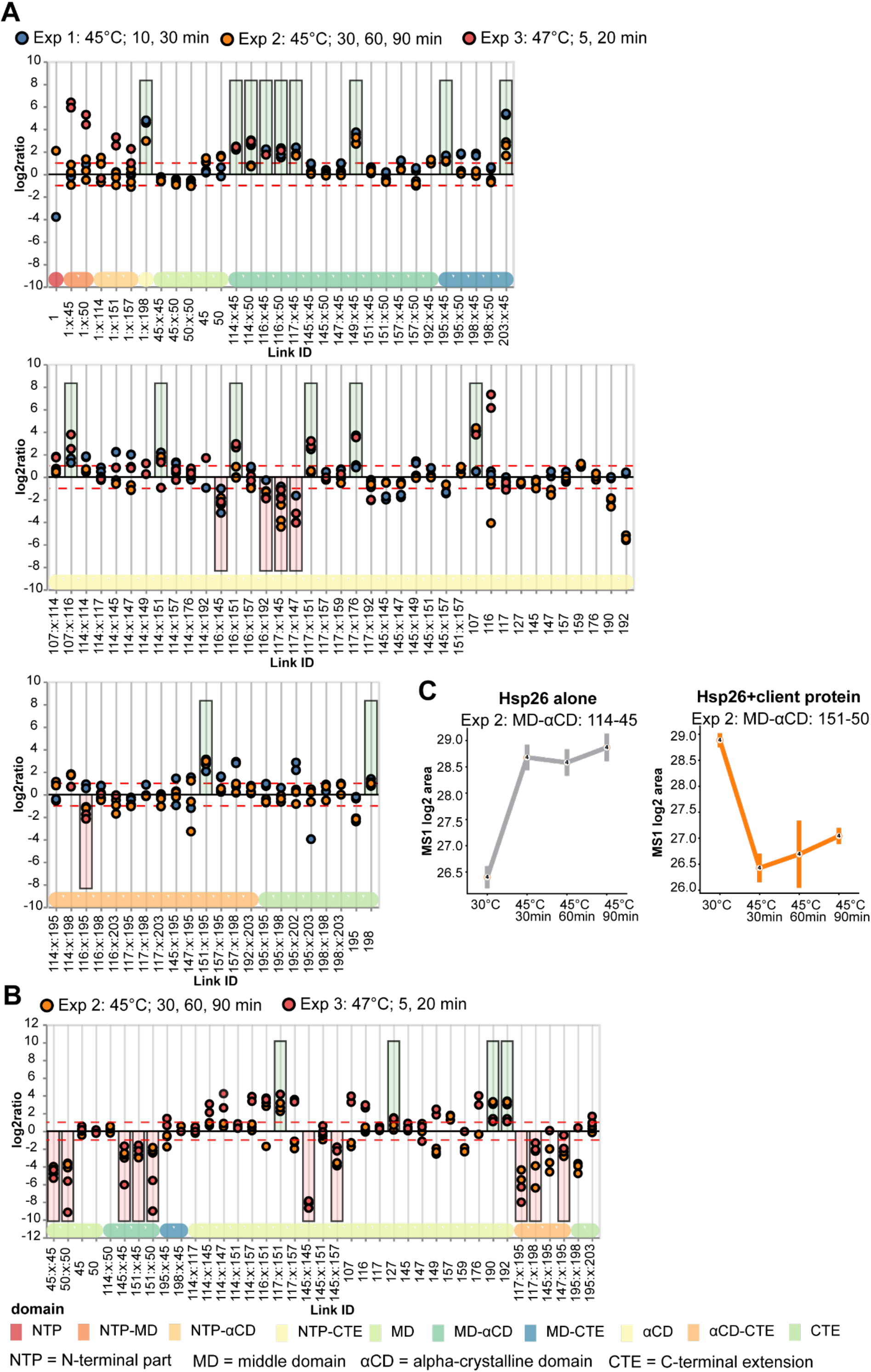
Time-resolved quantitative cross-linking mass spectrometry of Hsp26 in absence and presence of client protein. **(a)** Hsp26 in absence of a client protein. Plotted are the relative enrichment (heating vs 30°C) for each unique crosslinking site (y-axis) sorted according to the known domain structure within Hsp26 (x-axis);Circles in different shades of blue, orange and red)). Crosslinking sites that were consistently up – or downregulated two-fold or more (*see* methods and Supplementary Data 2 and 4 for details) in at least two out of three biological replicate sets and in addition contained no opposing regulation in any replicate set were considered significant and are highlighted with a green (enriched during heat stress) or red background rectangle (decreased during heat stress). All other changes in crosslinking abundances were considered insignificant and are shown on grey background. The significance threshold of two-fold enrichment is indicated as dashed red lines. **(Upper panel)** Unique cross-linking sites involving the NTD or MD; **(middle panel)** unique cross-linking sites within the alpha crystalline domain (αCD); **(lower panel)** unique cross-linking sites connecting the αCD to the C-terminal extension (CTE) and within the CTE. **(b)** Hsp26 in the presence of client (MDH). Experimental set-up and description as in (a), *see* methods for details. **(c)** Data for two selected cross-linking sites shown in (a) and (b). Shown are the extracted MS1 area (converted in log2 scale). Standard deviation is calculated based on cross-linking replicates as well as heavy and light cross-linked peptides (n = 4).

**Figure S5.**
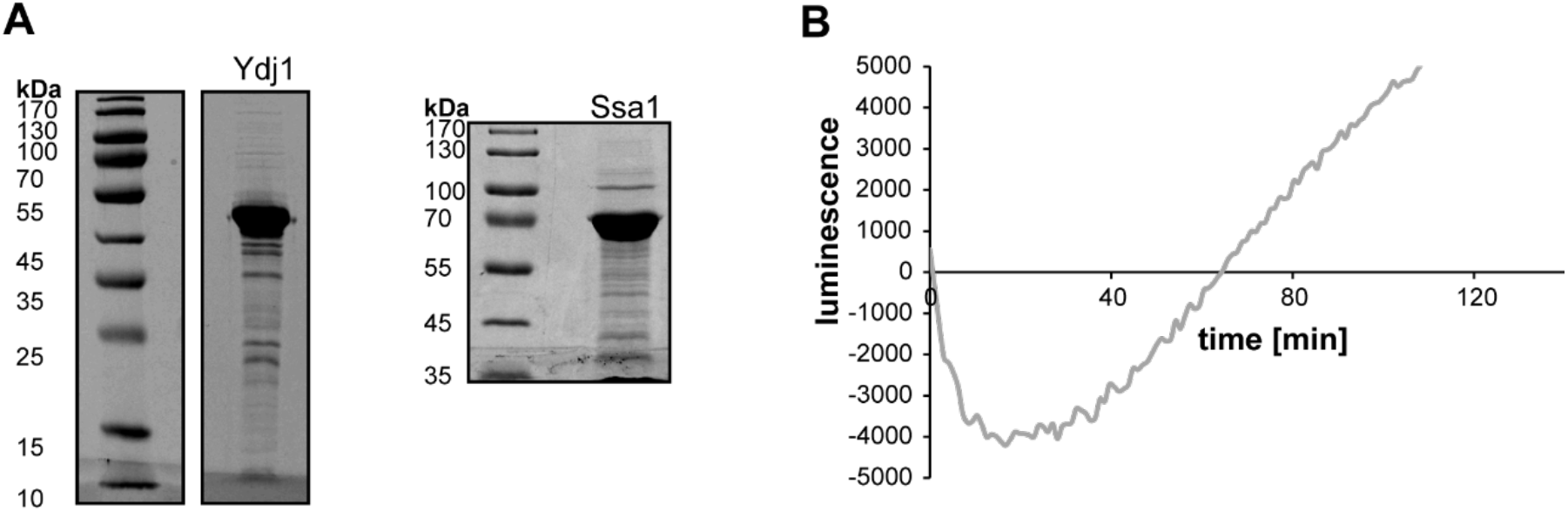
**(a)** SDS-gel of 10 µg of recombinantly expressed and purified Ydj1 and Ssa1. **(b)** Luciferase refolding assay in presence of Hsp26 and Ydj1/Ssa1. After heating of luciferase in the presence of Hsp26 and subsequent cool down for 30 min at 30°C, Ydj1 and Ssa1 were added, and luciferase activity was monitored continuously without addition of fresh substrate.

## Notes

### Competing Interest Statement

The authors have declared no competing interest.

